# Enhancement of Cell Adhesion by *Anaplasma phagocytophilum* Nucleolin-interacting Protein AFAP

**DOI:** 10.1101/2022.05.17.490364

**Authors:** Hongcheng Tang, Jiafeng Zhu, Shuyan Wu, Hua Niu

## Abstract

*Anaplasma phagocytophilum*, the aetiologic agent of human granulocytic anaplasmosis (HGA) is an obligate intracellular Gram-negative bacterium. During infection, *A. phagocytophilum* enhances the adhesion of neutrophils to infected endothelial cells. However the bacterial factors contributing to this phenomenon remain unknown. In this study, we characterized a type IV secretion system substrate of *A. phagocytophilum*, AFAP (an <u>a</u>ctin <u>f</u>ilament-associated *<u>A</u>naplasma phagocytophilum* <u>p</u>rotein), and found it enhanced cell adhesion. Tandem affinity purification combined with mass spectrometry identified host nucleolin as an AFAP-binding protein. Further study showed disruption of nucleolin by RNA interference and treatment of a nucleolin-binding DNA aptamer AS1411 attenuated AFAP-mediated cell adhesion. The characterization of AFAP with enhancement effect on cell adhesion and identification of its interaction partner may help understand the mechanism underlying *A. phagocytophilum*-promoting cell adhesion, facilitating elucidation of HGA pathogenesis.

**Highlights:**

- *Anaplasma phagocytophilum* AFAP localized to cell periphery.
- AFAP enhanced cell adhesion.
- AFAP interacted with host nucleolin.
- Disruption of nucleolin attenuated AFAP-mediated cell adhesion.

## 1. Introduction

*Anaplasma phagocytophilum*, a Gram-negative obligatory intracellular bacterium (rickettsia), causes the emerging tick-borne zoonosis called human granulocytic anaplasmosis (HGA), which is characterized with fever, malaise, headache, myalgia, arthralgia, leukopenia, thrombocytopenia, and elevations in serum hepatic aminotransferases (Bakken and Dumler, 2015). *A. phagocytophilum* infects midgut cells, hemocytes and salivary gland cells in ticks, and endothelial cells and neutrophils in mammalians (de la Fuente et al., 2016; Herron et al., 2005; Liu et al., 2011). It was proposed that after tick bite, endothelial cells serve as reservoirs for *A. phagocytophilum* and pass it on to neutrophils for dissemination and persistence in mammalian hosts (Herron et al., 2005; Wang et al., 2015). The efficient transmission of *A. phagocytophilum* from endothelial cells to neutrophils may rely on cell interaction. It was showed that during infection, *A. phagocytophilum* enhances the adhesion of neutrophils to infected endothelial cells (Herron et al., 2005; Wang et al., 2015). However bacterial factors contributing to the enhanced cell adhesion remain unknown.

Cell adhesion is mediated by adhesion molecules located on cell surface, which generally belong to five families: cadherins, integrins, selectins, immunoglobulin superfamily and others (Harjunpaa et al., 2019). These adhesion molecules engage their partners on other cells or extracellular matrix to form junctional contact or nonjunctional contact, promoting cells to move, communicate, differentiate, or assemble to form tissues and organs. Meanwhile with the linking to adhesion molecules via anchor proteins, cytoskeleton strengthens cell adhesion (Fuhrmann and Engler, 2015). There are several types of cell junctions, such as tight junctions, adherens junctions, desmosomes, and gap junctions in cell-cell adhesion, hemidesmosomes, focal adhesion, and podosomes in cell-matrix adhesion (Aghajanian et al., 2008; Block et al., 2008; Green and Jones, 1996; Linder and Kopp, 2005; Shimizu and Stopfer, 2013). These cell junctions play an important role in innate defence as exemplified by impermeability of epithelia, which blocks pathogen penetration, and exfoliation of mucosal epithelial cells, which inhibits bacterial colonization (Mulvey et al., 2000; Sansonetti, 2004). However, pathogens can manipulate cell adhesion to benefit their infection (Bonazzi and Cossart, 2011; Muenzner and Hauck, 2020). For instances, *Shigella flexneri* translocates its effector OspE into epithelial cells to enforce focal adhesion by interacting host integrin-linked kinase, thereby inhibiting cell detachment, and promoting bacterial colonization (Kim et al., 2009); *Listeria* secrets protein InlC into cytosol of infected epithelial cells, interacting with host protein Tuba to remodel cell-cell junctions, facilitating its spreading among host cells (Rajabian et al., 2009); HIV uses podosomes to invade macrophages (Li et al., 2021).

AFAP is a type IV secretion system effector of *Anaplasma phagocytophilum* and was found associated with host actin filaments, thereby named as an actin filament-associated *Anaplasma phagocytophilum* protein (Tang et al., 2020). We here found AFAP localized to cell periphery and enhanced cell adhesion. Tandem affinity purification combined with mass spectrometry identified host nucleolin as an interaction partner of AFAP. Further study showed disruption of nucleolin attenuated AFAP-mediated cell adhesion, indicating that AFAP enhanced cell adhesion likely through interaction with nucleolin. This study may help understand the mechanism underlying *A. phagocytophilum*-promoting cell adhesion, facilitating elucidation of HGA pathogenesis.

## 2. Materials and methods

### 2.1. Cell cultures and plasmids

HeLa cells and HEK293 cells were propagated in Dulbecco’s modified Eagle’s medium (DMEM) (Gibco) supplemented with 10% fetal bovine (Gibco) at 37°C and 5% CO_2_. Plasmids pAFAP-GFP and pLifeact-mCherry were constructed in previous study (Tang et al., 2020). Plasmid pEGFP-N1, which expresses GFP was from Clontech Laboratories. Plasmid pTAP-AFAP was constructed by cloning the gene fragment encoding AFAP-streptavidin-binding peptide-3xFLAG tag fusion protein (AFAP-SBP-3xFLAG, shorted as AFAP-SF) into mammalian expression vector pIRESpuro3 (Takara). SBP is a peptide with the length of 38 amino acids (Wilson et al., 2001); 3xFLAG is a epitope tag with the length of 22 amino acids (Zeghouf et al., 2004). The DNA fragment encoding SBP-3xFLAG was chemically synthesized and ligated to AFAP-coding DNA fragment, generated by PCR using pAFAP-GFP as template, to obtain AFAP-SF-coding DNA fragment. Plasmid pTAP, which expresses streptavidin-binding peptide-3xFLAG tag fusion protein (SBP-3xFLAG, shorted as SF) only was constructed by deletion of AFAP-coding sequence from pTAP-AFAP. The construction of plasmids pTAP-AFAP and pTAP was completed by GENEWIZ, Inc. (Suzhou, China).

### 2.2. Transfection, tandem affinity purification and coimmunoprecipitation

To determine subcellular localization of AFAP and actin filaments, plasmid pAFAP-GFP, or pEGFP-N1 was transfected into HeLa cells, together with pLifeact-mCherry by using Lipofectamine 3000 transfection reagent (Invitrogen), according to manufacturer’s instruction. To generate stable transfectants expressing AFAP-SF or SF, HEK293 cells were transfected with pTAP-AFAP or pTAP, respectively using Lipofectamine 3000 transfection reagent, followed by cultivation in complete DMEM medium containing 0.5 μg/mL puromycin (Thermo Fisher Scientific). To perform tandem affinity purification, HEK293 cells stably transfected with pTAP-AFAP or nontransfected HEK293 cells propagated in eight 150 mm^2^ cell culture dishes were lysed in lysis buffer (30 mM Tris-HCl, 150 mM NaCl, and 0.5% (v/v) nonidet-P40, pH 7.4) (1.2 mL/dish), supplemented with protease inhibitor cocktail and phosphatase inhibitors (APExBIO). The cell lysates were cleared by centrifugation at 12 000 *g* for 10 min at 4 °C, and the supernatants were subjected to incubation with streptavidin resin (GenScript) (50 μL/dish) for 2 h at 4 °C, followed by washing 4 times with washing buffer (30 mM Tris-HCl, 150 mM NaCl, and 0.1% (v/v) nonidet-P40, pH 7.4), supplemented with protease inhibitor cocktail and phosphatase inhibitors, and elution with elution buffer (30 mM Tris-HCl, 150 mM NaCl, and 4 mM biotin, pH 7.4). The eluates were further purified with magnetic beads conjugated with mouse anti-FLAG antibody (Bimake, Shanghai, China) (15 μL/dish). After incubation for 2 h at 4 °C, the beads were washed with washing buffer once and TBS (30 mM Tris-HCl, 150 mM NaCl, pH 7.4) twice, followed by elution with non-reducing 2 x SDS-PAGE sample loading buffer (100 mM Tris-Cl, 20% (v/v) glycerol, 4% (w/v) SDS, and 0.2% (w/v) bromophenol blue, pH 6.8). The eluates were subjected to SDS-PAGE and mass spectrometry for protein identification (BiotechPack Scientific, Beijing, China).

For coimmunoprecipitation, HEK293 cells stably transfected with pTAP-AFAP were lysed in immunoprecipitation buffer (IP) (50 mM HEPES, 150 mM KCl, 1 mM EDTA, 1.0% (v/v) Triton X-100, 10% (v/v) glycerol, pH 7.4) supplemented with protease inhibitor cocktail (APExBIO). After cell lysate was cleared by centrifugation at 16 000 *g* for 10 min at 4 °C, the supernatant was subjected to coimmunoprecipitation in a tube containing 2 μg rabbit polyclonal anti-FLAG antibody (Bioworld Technology, MN, USA), rabbit polyclonal anti-HA tag antibody (BBI Life Sciences, Shanghai, China), mouse monoclonal anti-nucleolin (NCL) antibody (D-6, Santa Cruz Biotechnology (SCBT)) or mouse normal IgG (SCBT). After incubation at 4 °C for 2 h, 20 μL protein A agarose (SCBT) was added into each tube. Protein A agarose resin was washed 4 times with IP buffer, and eluted by boiling for 5 min in 2 × SDS-PAGE sample loading buffer containing 200 mM DTT. The precipitates were probed in Western blot analysis using mouse monoclonal anti-nucleolin antibody (D-6) and rabbit polyclonal anti-FLAG antibody.

### 2.3. Immunofluorescence labeling

For determination of surface localization of nucleolin in AFAP-GFP-expressing HeLa cells, HeLa cells transfected with pAFAP-GFP for 3 d, were subjected to incubation at RT for 1 h with mouse monoclonal anti-nucleolin antibody (MS-3, SCBT) diluted in DMEM medium at 1:20. After washing with DMEM medium for 3 times, cells were fixed with 4% paraformaldehyde at RT for 20 min. Fixed cells were washed 3 times with 1 × PBS (137 mM NaCl, 2.7 mM KCl, 10 mM Na_2_HPO4, and 2 mM KH_2_PO4, pH 7.4) and blocked in 1 × PBS containing 0.8% BSA for 10 min, followed by incubation at 37 °C for 45 min with Alexa Fluor 555 conjugated goat anti-mouse antibody (Invitrogen). Cells were observed under Leica TCS SP8 confocal microscope (Leica).

### 2.4. RNA interference and treatment with AS1411 aptamer

For RNA interference, HEK293 cells stably transfected with pTAP-AFAP were transfected with siRNA against nucleolin (NCL) mRNA and control siRNA A (SCBT), using Lipofectamine RNAiMAX transfection reagent (Invitrogen), according to manufacturer’s instruction. Briefly, the day before transfection, 1 ×10^4^ cells were seeded into a well of 96-well plate. 1 pmol of each siRNA was added into HEK293 cells after formation of siRNA-lipid complex. The cells were further incubated for 3 d, followed by cell detachment assay, and Western blot analysis using mouse monoclonal anti-nucleolin antibody (D-6, SCBT) and mouse monoclonal anti-β-actin antibody (Beyotime Biotechnology, Shanghai, China).

AS1411 aptamer, which is a guanosine rich oligonucleotide, specifically targets cell surface nucleolin (Bates et al., 2009). AS1411 (5′-GGTGGTGGTGGTTGTGGTGGTGGTGG-3′) and its negative control CRO (cytosine rich oligonucleotide) (5′-CCTCCTCCTCCTTCTCCTCCTCCTCC-3′) were chemically synthesized, desalted (GENEWIZ), reconstituted in pure water at the concentration of 500 μM and filtered through 0.22 μm filter. AS1411 or CRO was added into 2 ×10^4^ HEK293 cells in a well of 96-well plate at final concentration of 2, 6 and 10 μM, respectively. After incubation for 4 d, the cells were subjected to cell detachment assay.

### 2.5. Cell detachment assay

Transfected HEK293 cells in 96-well plates were washed with prewarmed 1 × PBS after cultivation for 4 to 6 d, followed by incubation with 2 μM calcein-AM (Beyotime Biotechnology) in DMEM medium for 30 min. After DMEM medium containing calcein-AM was replaced with fresh DMEM medium, HEK293 cells were incubated for another 30 min. After washing twice with prewarmed 1 × PBS, the plates were sealed with plate cover and centrifuged upside down at 100 *g* for 5 min at 4 °C. Floating cells was carefully removed, and the wells were refilled with 1 × PBS. Cells were imaged and fluorescence intensity in wells was measured in a plate reader (BioTek) with excitation wavelength of 494 nm and emission wavelength of 514 nm before and after centrifugation.

### 2.6. Western blot analysis

HEK293 cells or immunoprecipitates in reducing 2 × SDS-PAGE sample loading buffer were subjected to SDS-PAGE with 10% polyacrylamide resolving gels. Proteins were transferred to nitrocellulose membranes using mini Trans-Blot cell (Bio-Rad), and the membranes were probed with primary antibodies including mouse monoclonal anti-nucleolin, rabbit anti-FLAG, or mouse monoclonal anti-β-actin at RT for 1 h. β-actin was used as an internal control for sample loading in RNA interference. After washing three times with 1 x PBS (10 min each time), the membranes were incubated with secondary antibody, peroxidase-conjugated goat anti-mouse IgG, or peroxidase-conjugated goat anti-rabbit IgG (KPL, Gaithersburg, MD) at RT for 1 h. The membranes were washed four times with 1 x PBS (10 min each time), and subjected to ECL chemiluminescence. The membranes were imaged by Tanon 4200 chemiluminescence imaging system (Tanon, Shanghai, China).

### 2.7. Statistical analysis

Three and more independent experiments were conducted and data were expressed as mean± SEM. Statistical significance was evaluated by Student’s *t*-test.

## 3. Results

### 3.1. AFAP dynamically changed its pattern and subcellular location in transfected cells

AFAP was previously found colocalized with the actin filaments at 24 h post-transfection in HeLa cells (Tang et al., 2020). However with the extended incubation time, a population of AFAP formed ring-like organization, which encircled punctate F-actin-rich structures, and another population of AFAP localized to cell periphery with actin filaments at 48 and 72 h posttransfection (Figure 1A and B). The punctate F-actin-rich structures and actin filaments were showed by label with F-actin-binding Lifeact, which is a yeast peptide with the length of 17 amino acids (Riedl et al., 2008). In contrast, no such punctate F-actin-rich structures were observed in GFP-expressing HeLa cells (Figure 1C). This result indicates AFAP dynamically changed its pattern and subcellular location in transfected cells over time. Furthermore, the stress fibers were clearly observed in GFP-expressing cells but rarely seen in AFAP-GFP-expressing cells at 72 h post-transfection (Figure 1), suggesting AFAP may cause the remodeling of actin cytoskeleton.

**Figure 1.**
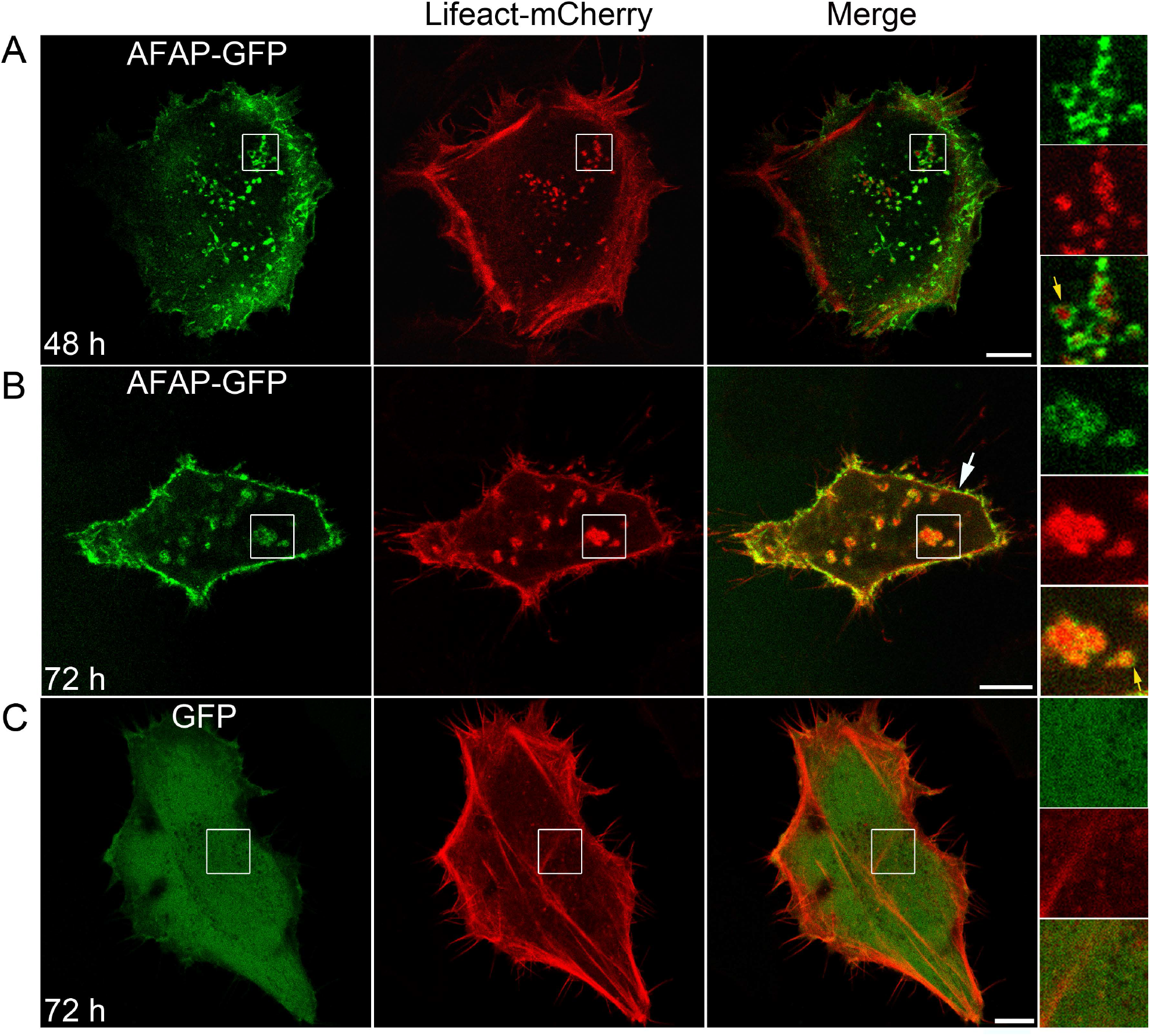
The dynamical changes of pattern and subcellular location of AFAP. HeLa cells were cotransfected with plasmids expressing AFAP-GFP and Lifeact-mCherry, followed by observation under confocal fluorescence microscope at 48 h post-transfection (A), and 72 h post-transfection (B). or cotransfected with plasmids expressing GFP and Lifeact-mCherry, followed by observation at 72 h post-transfection (C). Lifeact, an F-actin binding peptide. The boxed areas are magnified on the right. Yellow arrows indicate the structures with two-part architecture, in which F-actin cores were surrounded by AFAP rings. White arrow indicates cell periphery. Scale bars: 10 μm.

### 3.2 AFAP enhanced cell adhesion

punctate F-actin-rich structures encircled by rings are a prominent feature of podosomes (Linder and Kopp, 2005). Podosomes are adhesion structures formed in monocytic myeloid lineage and stimulated endothelial cells with two-part architecture, in which F-actin-rich cores are surrounded by rings composed of adhesion plaque proteins and integrins (Linder and Kopp, 2005; Veillat et al., 2015). In AFAP-expressing HeLa cells, the structures presented as F-actin-rich cores surrounded by AFAP rings were observed, morphologically similar to those of podosomes. Moreover, AFAP localized to cell periphery. Proteins at cell periphery participate in multiple cellular activities, such as cell adhesion, migration and signal transduction. Thus the effect of AFAP on these cellular activities, especially on cell adhesion was investigated.

Plasmid pTAP-AFAP, which expresses AFAP-streptavidin-binding peptide-3xFLAG tag fusion protein (AFAP-SF) was constructed for studying AFAP-mediated cellular activities, and identifying AFAP-interacting proteins in the following tandem affinity purification (Figure 2A). AFAP-SF is a protein with the length of 394 amino acids, including 38-amino-acid streptavidin-binding peptide and 22-amino-acid 3xFLAG tag (Figure 2B). When AFAP-SF was expressed in transfected cells, it showed expected size (Figure 2C). HEK293 cells have been employed for cell adhesion and migration studies (John Jayakumar et al., 2020; Kuang et al., 2009). While HEK293 cells are loosely adherent, which are easy to be dissociated from plastic surface by force, HEK293 cells stably expressing AFAP-SF were difficult to be dissociated from culture flask by pipetting in the routine cell passaging, especially with extended incubation time. Trypsin treatment had to be employed to completely dissociate the cells from culture flask when the cells were cultured for more than 6 d (data not shown). To determine the effect of AFAP on cell adhesion in a quantitative way, cell detachment assay was employed, which uses the centrifugation force to detach cell from plastic surface (Muenzner et al., 2005; Reyes and Garcia, 2003). No obvious cell dissociation was observed under light microscope after centrifugation in wells, in which HEK293 cells stably transfected with pTAP-AFAP (AFAP-SF) were cultured for 6 d, compared to HEK293 cells control stably transfected with pTAP (SF), which only had scattered cells in wells after centrifugation (Figure 2D). Meanwhile, 95.5±1.0% fluorescence intensity of calcein-AM was found remained in these AFAP-SF wells, compared to 7.5±1.7% fluorescence intensity in SF wells after centrifugation (Figure 2E). This result indicates AFAP enhanced cell adhesion.

**Figure 2.**
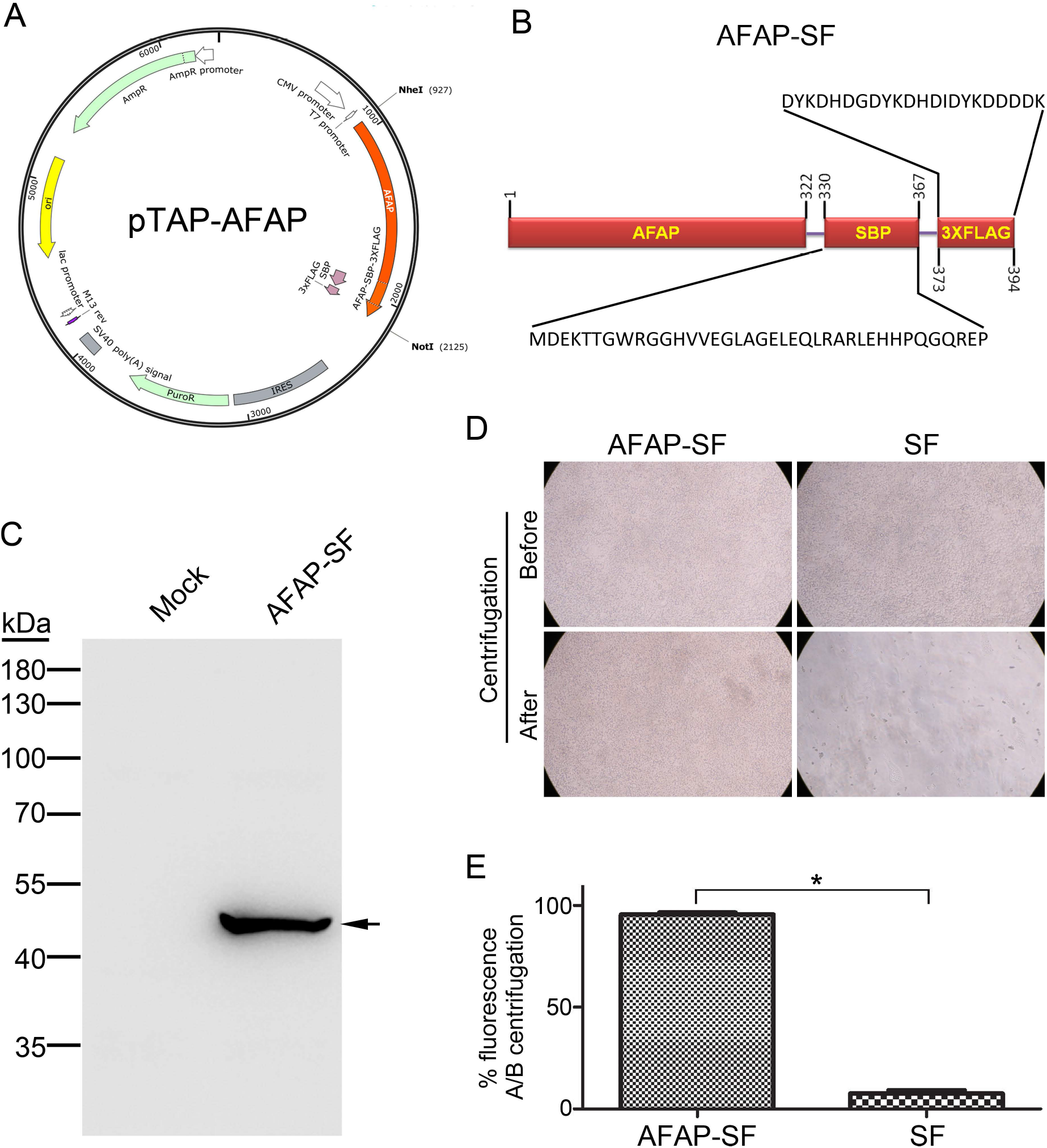
Construction of plasmid pTAP-AFAP, expression of AFAP-SF, and cell adhesion. (A) Construction of plasmid pTAP-AFAP expressing AFAP-SBP-3xFLAG (AFAP-SF). The gene encoding AFAP-SBP-3xFLAG was chemically synthesized, followed by cloning into plasmid pIRESpuro3 to generate pTAP-AFAP. (B) Diagram of AFAP-SBP-3xFLAG fusion protein (AFAP-SF). SBP (Streptavidin-binding peptide), a peptide with 38 amino acids in length. 3×FLAG, a peptide tag with 22 amino acids in length. The numbers indicate the start position and end position for each designated protein or peptide. (C) Western blot analysis for AFAP-SF in HEK293 cells transiently transfected with pTAP-AFAP (AFAP-SF). Mock: nontransfected HEK293 cells. Arrow: AFAP-SF band. (D) The cell images of HEK293 cells stably transfected with pTAP-AFAP (AFAP-SF) or pTAP (SF), were captured under light microscope (x100 magnification), before and after centrifugation. (E) Percentages of fluorescence intensity (fluorescence) of calcein-AM in wells after centrifugation/before centrifugation (A/B centrifugation) in cell detachment assay. AFAP-SF: HEK293 cells stably transfected with pTAP-AFAP. SF: HEK293 cells stably transfected with pTAP. *: Significant difference (*P* < 0.01) between groups indicated with lines by Student’s *t*-test.

### 3.3. Nucleolin was identified as an AFAP-interacting protein

AFAP-SF incorporated streptavidin-binding peptide and 3xFLAG, which were employed in tandem affinity purification to isolate AFAP-SF-interacting proteins. Besides AFAP-SF band, two extra bands with the size corresponding to 110 and 100 kDa, respectively were revealed by SDS-PAGE in eluate from HEK293 cells stably expressing AFAP-SF, compared to that from nontransfected HEK293 cells (Figure 3A). Although the band with the size of 100 kDa remained unidentified, the band with the size of 110 kDa was identified as nucleolin by mass spectrometry. To confirm the interaction between AFAP and nucleolin, coimmunoprecipitation was performed. Both anti-FLAG antibody and anti-nucleolin antibody pulled down the AFAP-SF and nucleolin (Figure 3B and C), indicating there is interaction between AFAP and nucleolin. Nucleolin is a multifunctional protein, participating in a variety of cell functions such as ribosome biogenesis, transcriptional regulation, cell proliferation, and apoptosis (Ginisty et al., 1999). While nucleolin is mainly found in the nucleus and cytoplasm, it has also been found on the cell surface (Christian et al., 2003; Said et al., 2002). Cell surface nucleolin binds a multitude of ligands, such as laminin-1 and L-selectin, mediating the cell-matrix adhesion and cell-cell adhesion (Goldson et al., 2020; Kibbey et al., 1995). AFAP also localized to cell periphery. To determine whether cell surface is the location, on which AFAP interacted with nucleolin, immunofluorescence labeling for cell surface nucleolin was performed in AFAP-GFP expressing HeLa cells. Cell surface nucleolin was found colocalized with AFAP-GFP (Figure 3D), indicating cell surface is a subcellular location for the interaction between AFAP and nucleolin.

**Figure 3.**
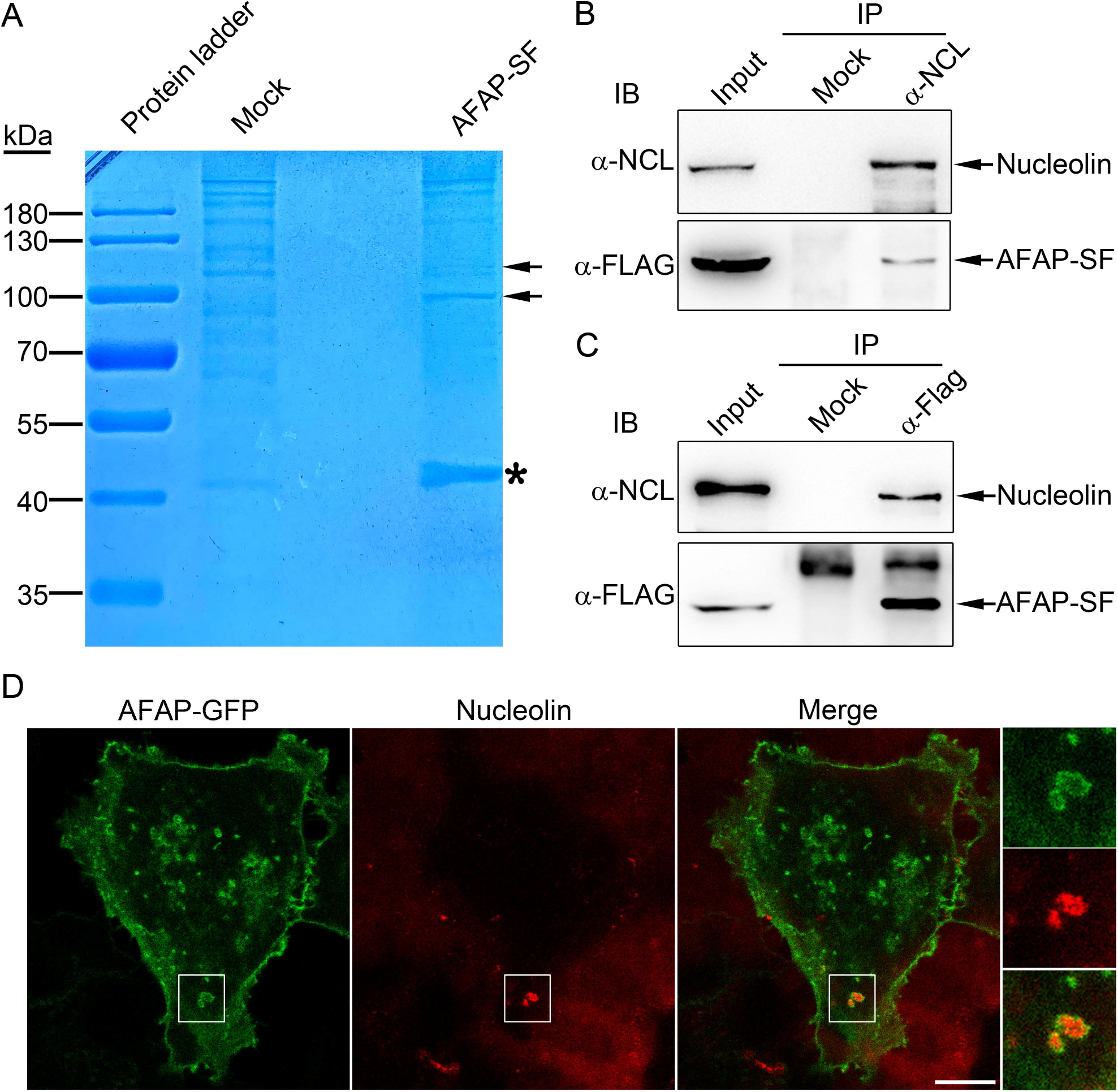
Identification of host nucleolin as an AFAP-interacting protein. (A) Tandem affinity purification of AFAP-interacting proteins and mass spectrometry. HEK293 cells stably transfected with pTAP-AFAP (AFAP-SF) and HEK293 cells (mock) were subjected to tandem affinity purification with streptavidin resin and magnetic beads conjugated with mouse anti-FLAG antibody after lysis, followed by SDS-PAGE. Asterisk: AFAP-SF band; Arrows: bands present in AFAP-SF lane but absent in mock lane. The bands indicated with arrows were subjected to mass spectrometry and the upper band was identified as nucleolin. (B-C) Coimmunoprecipitation assay for interaction between AFAP and nucleolin. The cell lysates form HEK293 cells stably transfected with pTAP-AFAP (AFAP-SF) were subjected to immunoprecipitated (IP) with mouse normal IgG (mock) (B), mouse monoclonal anti-nucleolin (α-NCL) (B), rabbit polyclonal anti-HA antibody (mock) (C), or rabbit polyclonal anti-FLAG antibody (α-FLAG) (C). Immunoprecipitates were immunoblotted (IB) with mouse monoclonal anti-nucleolin (α-NCL) and rabbit polyclonal anti-FLAG antibody (α-FLAG). (D) Colocalization assay for AFAP and nucleolin. HeLa cells were transfected for 3 d with plasmid expressing AFAP-GFP, followed by incubation with mouse monoclonal anti-nucleolin antibody and Alexa Fluor 555-conjugated goat anti-mouse IgG without permeabilization of cell membrane. The boxed areas with interest are magnified on the right. Scale bar: 10 μm.

### 3.4 The effect of AFAP on cell adhesion was attenuated by disruption of nucleolin

Since AFAP interacted with nucleolin, the effect of nucleolin on AFAP-mediated cell adhesion was determined. siRNA-mediated knockdown of nucleolin was performed. Compared to siRNA negative control, siRNA treatment for nucleolin reduced its expression and cell adhesion, as showed by Western blot analysis and cell detachment assay (Figure 4A and B). 79.2±2.7% fluorescence intensity of calcein-AM remained in wells of culture plate treated with siRNA negative control versus 52.1±2.0% fluorescence intensity of calcein-AM remained in wells of culture plate treated with nucleolin siRNA after centrifugation (Figure 4A). To corroborate this result, treatment with a DNA aptamer, AS1411 was performed. AS1411, a G-rich oligonucleotide, targets cell surface nucleolin and exerts its effect such as inhibition of cell proliferation, and blocking the binding of nucleolin to its ligands (Bates et al., 2009; Vester et al., 2021). HEK293 cells stably expressing AFAP-SF were treated with AS1411 or its negative control, CRO at 2, 6 and 10 μM for 4 d. Compared to CRO, cells treated with AS1411 at 2, 6 and 10 μM were less adherent to wells after centrifugation in cell detachment assay (67.1±4.8% vs. 35.8±7.6% at 2 μM; 67.8±2.3% vs. 29.6±3.7% at 6 μM; 74.2±2.6% vs. 32.1±4.9% at 10 μM) (Figure 4C), indicating AS1411 attenuated AFAP-mediated cell adhesion. Taken together, these results showed that disruption of nucleolin attenuated the enhancement effect of AFAP on cell adhesion, suggesting AFAP enhanced cell adhesion likely through interaction with nucleolin.

**Figure 4.**
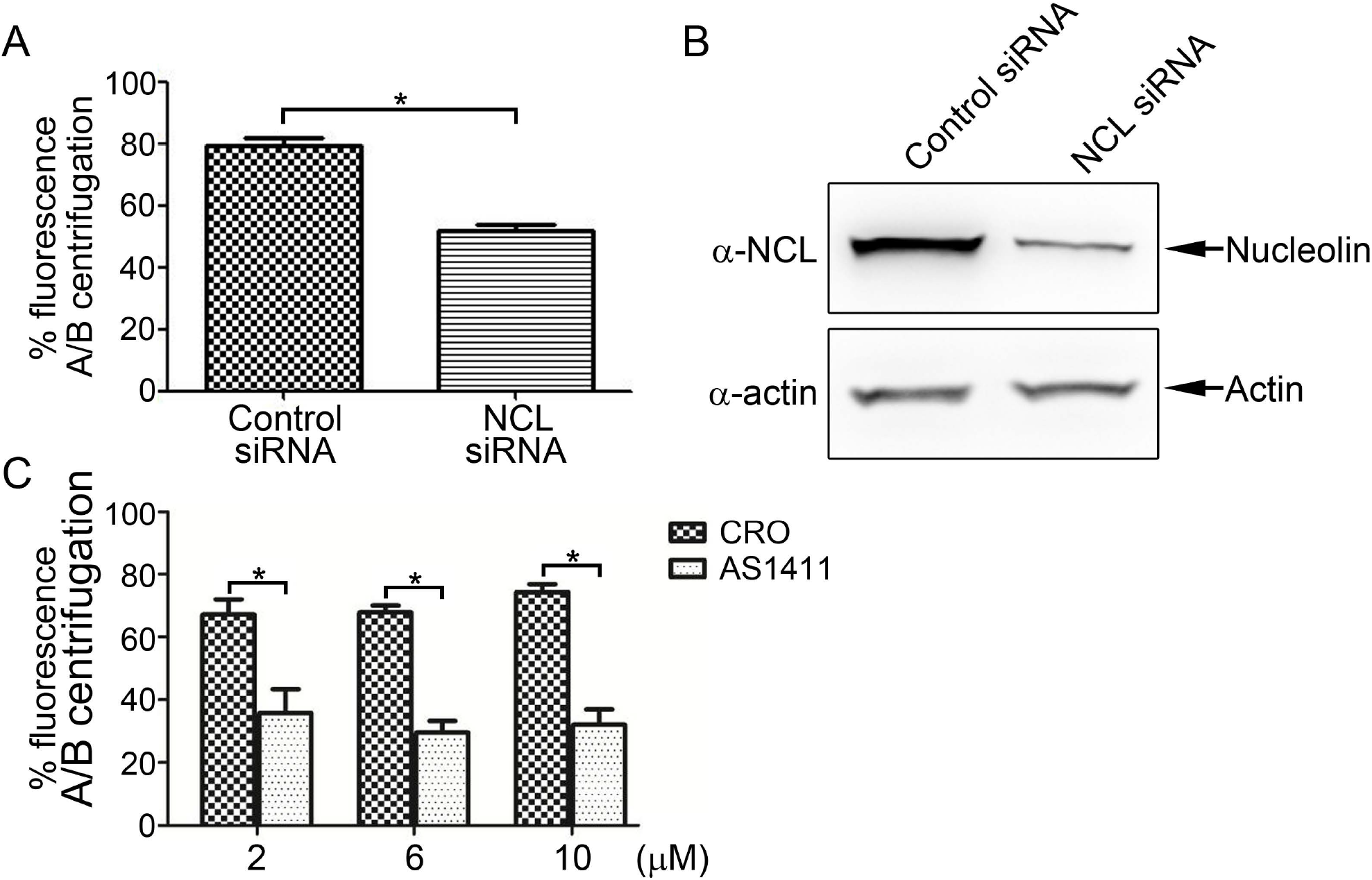
Disruption of nucleolin attenuated AFAP-mediated cell adhesion. (A) Treatment with nucleolin siRNA attenuated AFAP-mediated cell adhesion. HEK293 cells stably expressing AFAP-SF were transfected with control siRNA or nucleolin siRNA (NCL siRNA) for 3 d, followed by cell detachment assay. Percentages of fluorescence intensity (fluorescence) of calcein-AM in wells after centrifugation/before centrifugation (A/B centrifugation) were calculated. *: Significant difference (*P* < 0.01) between groups indicated with lines by Student’s *t*-test. (B) Treatment with nucleolin siRNA reduced nucleolin expression. HEK293 cells stably expressing AFAP-SF were transfected with control siRNA or nucleolin siRNA (NCL siRNA) for 3 d, followed by Western blot analysis using mouse anti-nucleolin (α-NCL) and anti-β-actin (α-actin) antibodies. Actin was used as an internal control to normalize sample loading amount. (C) Treatment with cell surface nucleolin-targeting aptamer AS1411 attenuated AFAP-mediated cell adhesion. HEK293 cells stably expressing AFAP-SF were treated with control DNA oligonucleotide (CRO) or aptamer AS1411 at designated concentrations for 4 d, followed by cell detachment assay. Percentages of fluorescence intensity (fluorescence) of calcein-AM in wells after centrifugation/before centrifugation (A/B centrifugation) were calculated. *: Significant difference (*P* < 0.05) between groups indicated with lines by Student’s *t*-test.

## 4. Discussion

*A. phagocytophilum* infection enhances the adhesion of neutrophils to infected endothelial cells (Herron et al., 2005; Wang et al., 2015). In this study, it was found that AFAP enhanced cell adhesion in transfected cells. Cell adhesion is mediated by adhesion molecules located on cell surface. AFAP was found localized to cell periphery at 48 and 72 h post-transfection, indicating it may participate in cell adhesion. Cell adhesion between cell and extracellular matrix is mainly mediated by junctional contact, such as focal adhesion and podosomes (Block et al., 2008; Linder and Kopp, 2005). Podosomes are characterized with two-part architecture, in which F-actin-rich cores are surrounded by rings composed of adhesion plaque proteins and integrins (Linder and Kopp, 2005; Veillat et al., 2015). In AFAP-expressing cells, the structures presented as F-actin-rich cores, surrounded by AFAP rings were observed. Whether these structures have the characteristics of podosomes and whether they are formed in infected cells need further investigation. Meanwhile we noticed that stress fibers in transfected cells were reduced at 72 h post-infection. It was showed that stress fibers were disassembled in infected tick cells (Sultana et al., 2010). Whether AFAP is contributed to this phenomenon in infected tick cells need further study.

In this study, host nucleolin was identified as an AFAP-interacting protein. The nucleolin-AFAP interaction location was likely on the cell surface, since nucleolin colocalized with AFAP in immunofluorescence labeling was stained without permeabilization of cell membranes. Although nucleolin is mainly found in the nucleus and cytoplasm, it has also been found on the cell surface (Christian et al., 2003; Said et al., 2002). Cell surface nucleolin is a receptor for multiple ligands, such as laminin-1, L-selectin, and virus surface proteins, mediating cell adhesion and pathogen entry (Goldson et al., 2020; Kibbey et al., 1995; Mastrangelo et al., 2021). L-selectin is abundantly expressed on circulating neutrophils and nucleolin is present on cell surface of endothelial cells (Christian et al., 2003; Rahman et al., 2021). It is worthwhile to determine the role of nucleolin in cell adhesion between infected endothelial cells and neutrophils. Furthermore it was reported that cell surface-expressing nucleolin is associated with actin filaments, likely through the interaction with myosin heavy chain 9 (Hovanessian et al., 2000; Huang et al., 2006). AFAP is an actin filament-associated protein, and found surrounding F-actin-rich structures. Thus nucleolin is likely colocalized with the F-actin-rich structures surrounded by AFAP.

## 5. Conclusions

In this study, we found AFAP localized to cell periphery, surrounded punctate F-actin-rich structures and enhanced cell adhesion. Host nucleolin was identified as an interacting protein of AFAP. Disruption of nucleolin attenuated AFAP-mediated cell adhesion, indicating that AFAP enhanced cell adhesion likely through interaction with nucleolin. This study may help understand the mechanism underlying *A. phagocytophilum-promoting* cell adhesion, facilitating elucidation of HGA pathogenesis.

## Author statement

**Hongcheng Tang:** Methodology, Formal analysis, Investigation. **Jiafeng Zhu:** Formal analysis. Shuyan Wu: Supervision. **Hua Niu:** Conceptualization, Methodology, Formal analysis, Writing-Review & Editing, Supervision, Project administration, Funding acquisition.

## Declaration of interest

The authors declare that they have no conflict of interest.

## Funding

This study was funded by the National Natural Science Foundation of China [grant number 31470276] and Guangxi Natural Science Foundation [grant number 2021GXNSFAA196007].

## Acknowledgments

The authors wish to thank Zhonglin Fu in the Institutes of translational Medicine at Soochow University for providing technical assistance in confocal microscopy.

## Notes

### Competing Interest Statement

The authors have declared no competing interest.

